# Estimating prevalence for limb-girdle muscular dystrophy based on public sequencing databases

**DOI:** 10.1101/502708

**Authors:** Wei Liu, Sander Pajusalu, Nicole J. Lake, Geyu Zhou, Nilah Ioannidis, Plavi Mittal, Nicholas E. Johnson, Conrad C. Weihl, Bradley A. Williams, Douglas E. Albrecht, Laura E. Rufibach, Monkol Lek

**Author notes:** Contributed equally to the work (Sander Pajusalu, Nicole J. Lake). To whom correspondence should be addressed: Monkol Lek, Department of Genetics, Yale School of Medicine, 300 Cedar Street, New Haven, CT 06520, USA.

## Abstract

**Purpose:** Limb Girdle Muscular Dystrophies (LGMD) are a genetically heterogeneous category of autosomal inherited muscle diseases. Many genes causing LGMD have been identified, and clinical trials are beginning for treatment of some genetic subtypes. However, even with the gene-level mechanisms known, it is still difficult to get a reliable and generalizable prevalence estimation for each subtype due to the limited amount of epidemiology data and the low incidence of LGMDs.

**Methods:** Taking advantage of recently published whole exome and genome sequencing data from the general population, we used a Bayesian method to develop a reliable disease prevalence estimator.

**Results:** This method was applied to nine recessive LGMD subtypes. The estimated disease prevalence calculated by this method were largely comparable to published estimates from epidemiological studies, however highlighted instances of possible under-diagnosis for LGMD2B and 2L.

**Conclusion:** The increasing size of aggregated population variant databases will allow for robust and reproducible prevalence estimates of recessive disease, which is critical for the strategic design and prioritization of clinical trials.

## Introduction

The limb girdle muscular dystrophies (LGMDs) are a heterogeneous group of diseases, causing pelvic and shoulder girdle muscle weakness and wasting. There are currently 32^1^ characterized subtypes with a diverse range of clinical phenotypes, which show variability in age of onset, rate of progression, specific muscle wasting patterns, and involvement of respiratory and cardiac muscles. The subtypes are broadly categorized by their pattern of inheritance as either dominant (LGMD1A-I) or recessive (LGMD2A-X) with the majority being recessive and can harbor either loss of function or missense pathogenic variants. The proteins encoded by LGMD disease genes have cellular functions including glycosylation, and muscle membrane integrity, maintenance and repair, which are a diverse range of mechanisms that when disrupted all result in muscle damage and degeneration.

Currently, an effective treatment does not exist for any LGMD subtype, however promising gene therapy clinical trials have commenced for LGMD2E and additional subtypes are set to commence in 2019-2020^2^. Disease prevalence information is critical to the planning and prioritization of these clinical trials. Historically, the prevalence of rare diseases has largely been estimated from epidemiological surveys and patient registries^3, 4, 5^. However, it can be difficult to achieve an accurate and meaningful prevalence estimate for rare genetic disorders through these traditional approaches. Many patients with rare disease experience a delayed or incorrect diagnosis, which can be more pronounced for late onset, slowly progressing diseases such as LGMD^6^, leading to under-estimation of prevalence. Differences in the diagnostic criteria used between studies, as well as changes to these over time, can make it difficult to directly compare estimates across studies. The specific population studies can also bias the prevalence estimate, and indeed the current published prevalence of LGMD subtypes can vary greatly between countries and even regions within countries^7^. The factors contributing to these regional differences include small sample size, founder mutations and consanguinity rates; all of which can lead to increased incidences of LGMD in those populations^8, 9, 10^. In addition, the resources and training available to each healthcare system can contribute to regional variability. Improved methods for quantifying the prevalence of rare genetic disorders such as LGMD are thus needed.

Using variants identified by large human exome and genome research studies as population references has greatly aided the filtering and interpretation of variants found in individuals with rare disease, and the study of known disease mutations in the general population^11^. The growth of these population genetic databases has enabled allele frequency data to be more widely used for estimating disease prevalence. However, there have been two main challenges with using allele frequencies from population reference databases to estimate prevalence. Firstly, the sample sizes can be insufficient to robustly estimate allele frequencies associated with rare diseases for which the majority of pathogenic variants are observed rarely in the general population. In addition, many databases have been inadequate for the estimation of disease prevalence in non-European populations. Although Bayesian methods for estimating disease prevalence have been developed and applied to allele frequency from large databases^12^, they currently do not incorporate separate prior distributions for each functional annotation (e.g. nonsense, missense, etc.).

In this study, we used publicly available population references to obtain a more reliable disease prevalence estimation for recessive LGMD (LGMD2). Previous epidemiology studies (**Table 1**) and approaches using population reference panels have been biased and would vary a lot across different reference databases when using allele frequencies based on one single observation. Although, overlapping variants from reference databases have similar allele frequencies for common variants (>0.5%), they may differ greatly at lower allele frequencies (<0.5%). In fact, 69% of European singletons within the Exome Sequencing Project (ESP) are not observed in the much larger ExAC data set^13^. To overcome this bias, we introduced a Bayesian method here to re-estimate allele frequencies, taking advantage of prior knowledge in the overall distributions of allele frequencies for different functional annotations (e.g. missense, frameshift, etc.). We developed a Bayesian framework to gain robust prevalence estimates with a confidence interval. By utilizing population reference panels ExAC and gnomAD, we simultaneously re-estimated allele frequencies for various functional annotations via a Bayesian method and then estimated disease prevalence assuming Hardy-Weinberg equilibrium. Our method uses the largest available population reference panels, which maximizes the capture of extremely rare variants. Furthermore, by categorizing variants based on their functional annotation, our method obtained improved estimates. Our approach can provide population-level disease prevalence estimation along with estimates for specific sub-populations like non-Finnish Europeans, and provides a generalizable and reproducible estimation compared with previous epidemiological approaches, which are sensitive to regional differences.

**Table 1:** Estimated prevalence (per million) in nine LGMD2 subtypes and published estimates in epidemiology studies.

| Subtype /gene | Population | Bayesian estimator (per million)+ 95% CI | Direct estimator (per million) | Estimates published in epidemiology studies (per million) |
| --- | --- | --- | --- | --- |
| 2A/CAP N3 | AFR | 27.0 (16.6, 39.1) | 26.5 | 9.47 in northeastern Italy<sup>17</sup><br>6.0 in northern England<sup>18</sup><br>4300 in a Mexican village<sup>10</sup><br>576 in a province of Spain<sup>36</sup><br>48 in the Reunion island<sup>37</sup> |
|  | ASJ | 0.02 (2.6e-5, 0.09) | 0.01 |  |
|  | EAS | 13.6 (4.7, 25.6) | 13.0 |  |
|  | EUR | 7.0 (5.4, 8.8) | 7.0 |  |
|  | FIN | 0.50 (0.12, 1.1) | 0.44 |  |
|  | NFE | 9.4 (7.1, 11.8) | 9.3 |  |
|  | All | 8.4 (6.8, 10.2) | 8.3 |  |
| 2B/DYS F | AFR | 34.2 (22.1, 48.2) | 16.0 | 1.3 in northern England<sup>6</sup> |
|  | ASJ | 0.06 (1.5e-4, 0.24) | 0.04 |  |
|  | EAS | 14.7 (3.5, 31.2) | 11.2 |  |
|  | EUR | 4.4 (2.8, 6.2) | 3.3 |  |
|  | FIN | 0.2 (0.03, 0.6) | 0.1 |  |
|  | NFE | 6.0 (3.7, 8.5) | 4.6 |  |
|  | All | 7.5 (5.7, 9.4) | 7.4 |  |
| 2C/SG CG | AFR | 0.4 (0.05, 1.0) | 0.4 | 1.3 in northern England<sup>18</sup><br>1.72 in northeastern Italy<sup>38</sup><br>70 in Tunisia<sup>39</sup><br>48.8 in Moroccan population<sup>40</sup><br>1.8 in Japan<sup>41</sup> |
|  | ASJ | 0.07 (1.7e-4, 0.3) | 0.05 |  |
|  | EAS | 0.02 (5e-05, 0.08) | 0.01 |  |
|  | EUR | 0.13 (0.04, 0.20) | 0.12 |  |
|  | FIN | 0.01 (3.2e-05, 0.05) | 0.01 |  |
|  | NFE | 0.17 (0.05, 0.3) | 0.16 |  |
|  | All | 0.12 (0.05, 0.20) | 0.11 |  |
| 2D/SG CA | AFR | 18.3 (10.1, 28.0) | 18.0 | 0.7 in northeastern Italy<sup>17</sup><br>3.02 in northeastern Italy<sup>38</sup> |
|  | ASJ | 3.5 (1.0, 7.2) | 3.3 |  |
|  | EAS | 0.3 (0.02, 0.7) | 0.2 |  |
|  | EUR | 3.9 (3.0, 5.0) | 3.9 |  |
|  | FIN | 5.9 (3.2, 9.1) | 5.7 |  |
|  | NFE | 3.6 (2.6, 4.7) | 3.5 |  |
|  | All | 3.4 (2.6, 4.2) | 3.3 |  |
| 2E/SGC B | AFR | 2.3 (0.4, 5.2) | 2.1 | 0.7 in northeastern Italy<sup>17</sup><br>0.86 in northeastern Italy<sup>39</sup> |
|  | ASJ | 0.6 (0.03, 1.5) | 0.5 |  |
|  | EAS | 0.14 (0.002, 0.4) | 0.11 |  |
|  | EUR | 0.6 (0.3, 1) | 0.6 |  |
|  | FIN | 0.07 (0.002, 0.2) | 0.06 |  |
|  | NFE | 0.8 (0.4, 1.3) | 0.8 |  |
|  | All | 0.80 (0.42, 1.26) | 0.78 |  |
| 2F/SGC<br>D | AFR | 0.2 (0.002, 0.5) | 0.12 | Not available |
|  | ASJ | 0 (0, 0) | 0 |  |
|  | EAS | 0.9 (0.001, 3.7) | 0.5 |  |
|  | EUR | 0.06 (0.004, 0.15) | 0.05 |  |
|  | FIN | 0.6 (0.002, 2) | 0.4 |  |
|  | NFE | 0.02 (0.004, 0.04) | 0.02 |  |
|  | All | 0.07 (0.01, 0.15) | 0.06 |  |
| 2G/TCA<br>P | AFR | 0.05 (4e-04, 0.2) | 0.04 | Not available |
|  | ASJ | 0 (0, 0) | 0 |  |
|  | EAS | 1.2 (0.4, 2.3) | 1.1 |  |
|  | EUR | 0.02 (0.004, 0.03) | 0.01 |  |
|  | FIN | 0.0001 (1.2e-07, 6e-4) | 0 |  |
|  | NFE | 0.02 (0.006, 0.05) | 0.02 |  |
|  | All | 0.040 (0.020, 0.063) | 0.039 |  |
| 2I/FKR<br>P | AFR | 1.5 (0.2, 3.4) | 1.4 | 4.3 in northeastern Italy<sup>17</sup> |
|  | ASJ | 0.03 (4.3e-05, 0.1) | 0.02 |  |
|  | EAS | 4.2 (1.5, 7.9) | 4.1 |  |
|  | EUR | 8.4 (5.7, 11.4) | 8.3 |  |
|  | FIN | 7.7 (1.8, 16.3) | 7.1 |  |
|  | NFE | 8.5 (5.7, 11.7) | 8.4 |  |
|  | All | 4.52 (3.20, 6.00) | 4.48 |  |
| 2L/ANO<br>5 | AFR | 4.8 (2.2, 8) | 4.6 | 2.7 in northern England<sup>8</sup><br>20 in Finland<sup>42</sup><br>10 in Denmark<sup>43</sup> |
|  | ASJ | 3.0 (0.8, 6.3) | 2.9 |  |
|  | EAS | 0.5 (0.09, 1.1) | 0.5 |  |
|  | EUR | 28.5 (24, 33.2) | 28.3 |  |
|  | FIN | 34.4 (35.8, 25.2) | 35.3 |  |
|  | NFE | 27.3 (22.4, 32.5) | 27.1 |  |
|  | All | 17.6 (15.2, 20.2) | 17.5 |  |

The table shows the comparison results between prevalence of LGMD2 subtypes estimated by our method (“Bayesian estimator”), by using the allele frequencies provided in gnomAD directly (“Direct estimator”) and from epidemiology studies. CI stands for confidence interval. The table also shows the population-stratification results for estimated LGMD2s prevalence. “AFR”: African/African America; “ASJ”: Ashkenazi
Jewish; "EAS": East Asian; "EUR": European; "FIN": Finnish; "NFE": Non-Finnish European; All: for mixed population in gnomAD

Utilizing this method, our estimates for LGMD2 subtypes are mainly consistent with published prevalence estimates along with a confidence interval. We also applied the method to another public genetic database (BRAVO) and found that prevalence estimates for 6 out of 9 LGMD2 subtypes using BRAVO were within the 95% prevalence confidence intervals estimated using gnomAD. Lastly, we applied our method to well characterized monogenic diseases Tay Sachs disease, sickle cell anemia and cystic fibrosis to validate our prevalence estimates. Overall, we provide a generalizable and robust framework to estimate disease prevalence for LGMD2 subtypes that can be easily adapted for other autosomal recessive diseases.

## Results

### Prevalence estimates in LGMD2 subtypes are comparable to published values

The recessive LGMDs (LGMD2) are autosomal recessive diseases, which can be caused by pathogenic variants in at least 24 genes^1^. We applied our Bayesian method to nine subtypes of LGMD2 from 2A to 2L (**Table 1**). The gnomAD dataset was used to identify putative and reported pathogenic variants in each disease gene. In brief, a variant in gnomAD was defined as pathogenic if it was either a loss of function variant present at <0.05% allele frequency, or annotated as pathogenic in the ClinVar or EGL databases. Using these variants, disease prevalence was then estimated for each gene (see Methods), along with 95% confidence intervals (**Table 1**), using the assumption that homozygous and compound heterozygous variants have the same penetrance.

The disease prevalence estimates calculated by our Bayesian method were generally consistent with published prevalence estimates from epidemiological studies (**Table 1**), in particular for LGMD2A, LGMD2E and LGMD2I. For other subtypes, our method produced a higher estimated prevalence, including LGMD2B, LGMD2D and LGMD2L. These differences can be partly explained by the under diagnosis of these late-onset or slowly progressive LGMD subtypes^14, 15^. In contrast, our disease prevalence estimation for subtype LGMD2C (0.12 per million) was notably lower than the lowest published value (1.3 per million). Genetics differences across regions would also contribute to discrepancies between our results and published estimators, since most epidemiology studies have been conducted in small regions, while the databases we are using include individuals with diverse genetic backgrounds. Lastly, no comparison could be made for LGMD2F and LGMD2G as there are no published prevalence estimates.

Next, we applied our method to another genetics database, BRAVO, to estimate prevalence for the same nine LGMD2 subtypes. When applied to a different database, our method provided more robust results compared with direct prevalence estimation (see **Methods**) using genetic data. Prevalence estimates for 6 out of 9 subtypes estimated in BRAVO fell in the 95% confidence intervals (CIs) estimated from the gnomAD data. The other three subtypes (2A, 2D and 2I) had an estimated prevalence close to the lower bounds of the corresponding 95% CI (**Table 2**). The differences of applying the same method (either our Bayesian method or the direct way, see **Methods**) in two different databases are much larger than that of using two methods in the same dataset, indicating the larger influence is the database used versus the method. The large differences in results from different databases can be partly explained by the sampling biases and limited sample size in each genetics dataset.

**Table 2:** Estimated disease prevalence (per million) in gnomAD and BRAVO for nine LGMD2s

| Subtype/gene | Direct in BRAVO, per million | Direct in gnomAD, per million | Bayesian in BRAVO, per million | Bayesian in gnomAD (95% CI), per million |
| --- | --- | --- | --- | --- |
| 2A/<br>CAPN3 | 6.2 | 8.3 | 6.3 | 8.4 (6.8, 10.2) |
| 2B/<br>DYSF | 5.9 | 7.4 | 6.0 | 7.5 (5.7, 9.4) |
| 2C/<br>SGCG | 0.09 | 0.1 | 0.09 | 0.1 (0.05, 0.2) |
| 2D/<br>SGCA | 2.0 | 3.3 | 2.0 | 3.4 (2.6, 4.2) |
| 2E/<br>SGCB | 0.5 | 0.8 | 0.5 | 0.8 (0.4, 1.3) |
| 2F/<br>SGCD | 0.03 | 0.06 | 0.04 | 0.07 (0.01, 0.15) |
| 2G/<br>TCAP | 0.02 | 0.04 | 0.02 | 0.04 (0.02, 0.06) |
| 2I/FKRP | 3.1 | 4.5 | 3.1 | 4.5 (3.2, 6.0) |
| 2L/<br>ANO5 | 15.3 | 17.5 | 15.4 | 17.6 (15.2, 20.2) |

This table lists the result of applying the direct method and our Bayesian method to estimate prevalence of 9 subtypes of LGMD2.

### Including rare missense variants currently not reported as pathogenic increases prevalence estimates

The above prevalence estimates are limited to reported pathogenic and rare loss of function variants found in gnomAD and do not account for other unreported missense pathogenic variants that may be in gnomAD. When we included all rare missense variants (AF < 0.05%), not surprisingly the prevalence estimates increased dramatically (**Table 3**) compared to the results indicated above. This increased prevalence was proportional to the coding length of the gene, as larger genes will accumulate more rare variants by random chance. The *DYSF* gene (80.4 kb) is much longer than *CAPN3* (29.2 kb) and therefore the fold change of prevalence estimates for including all rare missense variants versus only annotated pathogenic ones is larger (168 for *DYSF* versus 16.8 for *CAPN3*).

**Table 3:** Prevalence estimated (per million) including more predicted pathogenic variants

| Subtype/Gene | Annotated pathogenic and loss-function variants (same as Column 5 in table 2) (per million) | All rare missense included (per million) | CADD 20 cut-off included (per million) | CADD 30 cut-off included (per million) | Exon length (Kb) | Ratio of CADD 30 cut-off estimates/estimates in 2<sup>nd</sup> column |
| --- | --- | --- | --- | --- | --- | --- |
| 2A/<br>CAPN3 | 8.4<br>(6.8, 10.2) | 138<br>(125, 153) | 99<br>(91, 108) | 22<br>(19, 25) | 29.2 | 2.6 |
| 2B/<br>DYSF | 7.5<br>(5.7, 9.4) | 1260<br>(1190, 1320) | 620<br>(590, 650) | 105<br>(96, 115) | 80.4 | 14 |
| 2C/<br>SGCG | 0.1<br>(0.05, 0.2) | 19.6<br>(16.8, 22.6) | 7.8<br>(6.6, 9.1) | 0.9<br>(0.6, 1.2) | 1.6 | 9 |
| 2D/<br>SGCA | 3.4<br>(2.6, 4.2) | 43<br>(38, 49) | 18<br>(15, 20) | 6.1<br>(5.0, 7.3) | 9.0 | 1.8 |
| 2E/<br>SGCB | 0.8<br>(0.4, 1.3) | 28<br>(23, 33) | 8.0<br>(6.4, 9.7) | 1.1<br>(0.6, 1.7) | 5.7 | 1.4 |
| 2F/<br>SGCD | 0.07<br>(0.01, 0.15) | 10<br>(8, 12) | 4.4<br>(3.6, 5.4) | 0.3<br>(0.2, 0.5) | 13.5 | 4.3 |
| 2G/<br>TCAP | 0.04<br>(0.02, 0.06) | 4.7<br>(3.8, 5.8) | 4.2<br>(3.5, 4.9) | 0.26<br>(0.17, 0.35) | 2.7 | 6.5 |
| 2I/<br>FKRP | 4.5<br>(3.2, 6.0) | 65<br>(56, 75) | 32<br>(27, 37) | 6.7<br>(5.0, 8.5) | 19.7 | 1.5 |
| 2L/<br>ANO5 | 17.6<br>(15.2, 20.2) | 133<br>(121, 147) | 66<br>(60, 72) | 25<br>(22, 28) | 14.0 | 1.4 |

The table shows disease prevalence estimated by our Bayesian method when including computationally predicted pathogenic variants filtered by CADD score cut-offs. Numbers listed in brackets are the corresponding 95% confidence intervals.

This analysis assumes all rare missense variants are pathogenic, which is likely not the case. We then applied the Combined Annotation Dependent Depletion (CADD)^16^ method to classify the pathogenicity of rare missense variants. The CADD Phred-scaled cut-off scores of 20 and 30 were used to define pathogenicity, which respectively represent the top 1% and 0.1% of most deleterious substitutions predicted by the CADD method, ie the higher the score the more likely a variant will be pathogenic. The published prevalence estimates still fell outside of the 95% CI calculated when missense variants with a cut-off score of 20 were included, while the more stringent cut-off score of 30 produced closer estimates (see **Table 3**). For example, with LGMD2E, the estimated prevalence using a cut-off score of 30 is 1.1 per million, similar to the published 0.7 or 0.86 per million and is within the 95% CI (0.4 to 1.3) estimated when only considering rare loss-of-function variants and variants annotated pathogenic in ClinVar or EGL. These results show that improved pathogenicity predictions methods are required to improve disease prevalence estimates.

### Comparison with epidemiological results in population stratified analysis

The majority of epidemiological studies estimating disease prevalence have been conducted in small regions, leading to varying results across publications. LGMD2A serves as an example, where the estimates vary greatly in two small regions of Italy (6.1 and 16.5 per million)^17^. Due to the majority of the published estimates being from European populations, we limited our analysis to the sub-populations of European (EUR), Finnish (FIN) and non-Finnish European (NFE) here, results of more sub-populations are shown in **Table 1**.

After applying population stratification, estimated prevalence is more comparable with previously published results (see **Table 1**). For LGMD2A, the prevalence was estimated at 9.4 per million (95% CI: 7.1-11.8 per million) in the NFE population, matching the published value of 9.4 per million in northeastern Italy. However, after population stratification, the prevalence estimations for some subtypes diverged further from the published values. For LGMD2L, compared with the estimator (17.6 per million) in a mixed population, the estimator (27.3 per million) in the NFE population is even higher than the published prevalence (2.6 per million) in northern England^18^. The much higher result could be caused by the elevated allele frequency of a founder mutation, *ANO5* NM_213599.2:c.191dupA^8^, in the NFE population (0.21%) compared with the allele frequency in the mixed population (0.11%) in gnomAD. For subtypes only common in certain populations, the stratification can provide a more precise prevalence estimate. Take 2G for example, the prevalence estimated in East Asians (EAS) is about 1.2 per million while less than 0.05 per million in other populations (**Table 1**). This result suggests that varied genetic backgrounds can lead to population differences in disease prevalence estimates, which can be shown in results from both epidemiological studies and genetic databases.

### Inclusion of CADD in the prior and unseen pathogenic variants

We next performed two additional analyses specifically facilitated by using a Bayesian framework. CADD scores were used to categorize variants in combination with functional categories for updating allele frequencies of pathogenic variants (see **Methods**). After categorizing variants into smaller specific groups, disease prevalence were re-estimated for LGMD2 subtypes (**Supplementary Table 2**). Compared with results in **Table 1**, prevalence estimates were overall very similar with only small changes in sub-populations.

There are a number of reported pathogenic variants that were not observed in gnomAD (**Supplementary Table 1**) due to sampling and being ultra-rare variants. Using a Bayesian framework these variants can be included in the prevalence estimates resulting in slightly higher estimates in the sub-populations (**Supplementary Table 3**). Furthermore, this can provide a non-zero estimate and confidence intervals in instances where no pathogenic variants are observed in the sub-population (e.g. LGMD2G prevalence in Ashkenazi Jewish).

### Estimating prevalence in well studied diseases

To further confirm the reliability of our results, we also applied our method to three non-neuromuscular diseases; sickle cell disease, cystic fibrosis and Tay-Sachs disease, and estimated their prevalence in the sub-population where they were sourced. Pathogenic variants for the corresponding disease genes *HBB*, *CFTR and HEXA* were extracted from gnomAD and our Bayesian method was used to calculate the posterior allele frequency distributions and an estimate of disease prevalence.

Beta hemoglobinopathies including sickle cell disease and beta thalassemia are recessive blood disorders caused by mutations in HBB. Due to the p.Glu6Val missense mutation in HBB, sickle cell disease is prevalent in Africa and among people of African ancestry. The published prevalence is 2740 per million (one out of every 365) African-Americans^19^. Our estimated prevalence in African or African-Americans is 3490 per million (95% CI: 3140 - 3853 per million). For people with sickle cell disease, at least one of their two HBB alleles should have a specific variant (Hb S variant)^20, 21^, otherwise the patients would be diagnosed with other disease like beta thalassemia. Therefore, the estimated prevalence here includes the prevalence for other disease and therefore is higher than published figure for sickle cell anemia.

Cystic fibrosis is a recessive disorder caused by mutations in *CFTR*. The published mean prevalence in European populations is 74 per million across 27 European Union countries. Applying our method to the NFE population, the estimated prevalence is 365 per million (95% CI: 338 - 393 per million). Since our method does not consider time of onset of diseases, our prevalence estimates for life-shortening diseases such as cystic fibrosis will be closer to birth prevalence (incidence) than to population prevalence. After adjusting the published prevalence of 74 per million using the calibrating formula proposed by Farrell (incidence = 0.00019 + 0.00016 × prevalence per 10,000)^22^, an incidence of 308 per million in Europe was predicted, which is closer to our estimated prevalence of 365 per million.

Tay-Sachs disease is a neurological disorder caused by recessive mutations in the *HEXA* gene. The disease is prevalent in the Ashkenazi Jewish population due to several founder mutations including NM 000520.5:1274_1277dup and c.1421+1G>C^23^. Using the allele frequencies from the Ashkenazi Jewish sub-population, we estimate a prevalence of 197 per million, with a 95% CI 137 to 265 per million. The published 286 per million birth incidence figure^24^ is close to the upper bound of our estimation.

Overall, our prevalence estimates for these three diseases are similar to published figures, indicating that our method is reliable across multiple autosomal recessive diseases.

## Discussion

Through the application of a Bayesian method to large publicly available genetic databases, we have determined robust prevalence estimations for LGMD2 subtypes that are consistent with published figures from epidemiological studies. By applying our method of calculating prevalence to another genetics database, BRAVO, the robustness of the method was confirmed since most prevalence estimates from BRAVO were within our estimated confidence intervals using gnomAD. For further evaluation, we estimated prevalence for three non-muscular diseases using the method and generated similar values to published results.

Building upon a previous Bayesian prevalence estimation method^12^, we estimated LGMD2 prevalence by simultaneously considering more than one mutation using much larger databases, which mitigates underestimation of disease prevalence. We also considered functional annotation when updating allele frequency for each variant. Utilization of the largest genetic databases available also made our estimation more reliable, since databases with insufficient sample size would lead to increased absence of rare pathogenic variants.

Similar to other disease prevalence estimation methods based on genetic data, our estimation has strong assumptions that may affect our results. First, we assumed pathogenic variants observed in compound heterozygous and homozygous states have the same severity, which may result in differences compare to published values. For example, the c.191dupA founder mutation in *ANO5* is observed as homozygous in one individual in gnomAD suggesting later onset and/or a much milder muscle phenotype associated with being homozygous for this variant. Second, pathogenic variants are assumed to be independent of each other and therefore this method does not account for rare variants that occur on the same haplotype (i.e. linkage disequilibrium). Third, the analysis is limited to single nucleotide variants (SNVs) and small insertions and deletions. Large duplications and deletions account for some of the pathogenic variants discovered in neuromuscular disease genes with some having higher frequencies due to founder effects such as the exon 55 deletion in NEB^25^ associated with autosomal recessive Nemaline myopathy. Furthermore, we assume that all pathogenic variants for a subtype have been identified in the database we used here, which is likely not true (**Supplementary Table 1**), and may lead to an under-estimate of disease prevalence. Conversely, the current analysis does not take into account the situation where multiple disorders are caused by the variants in the same genes. For example, in the case of FKRP, the prevalence estimate includes both LGMD and Walker-Warburg syndrome variants^26, 27^, and thus can overestimate the LGMD prevalence. As mutation databases become more comprehensive, this information can be accurately extracted to mitigate this over estimation. Lastly, we have only estimated prevalence in this study for recessive LGMD2 disorders where compound heterozygous or homozygous mutations cause disease. Recently, several heterozygous mutations in genes associated with LGMD2 subtypes have been identified that can act dominantly, such as a 21-bp deletion in *CAPN3*^1^. This method we developed here is limited however for the estimation of dominant LGMD prevalence since dominant mutations are expected to be largely absent from population databases ExAC and gnomAD, while any present may be further complicated by reduced penetrance.

In contrast to published prevalence estimates from epidemiology studies, our results based on allele frequencies obtained from population reference databases are not impacted by public policy or health system specific to countries or regions. However, our results are also affected by different genetic backgrounds across regions (LGMD2L: 17.63 per million in global population and 27.33 per million in non-Finnish European population). Additionally, differences in sample sizes of various sub-populations in the genetic database used would also affect the identification of causal variants. Although the sample size of the database used here is the largest available, some rare pathogenic variants are likely to still be missing due to an insufficient sample size, which further leads to underestimation of prevalence, especially for rarer subtypes. The underestimated prevalence of LGMD2C (0.12 per million compared with 1.3 per million) may be caused in particular by the absence of various pathogenic variants in the database used. The Bayesian framework allowed for reported pathogenic variants unseen in gnomAD to be included in the prevalence estimates, however it did not result in much difference except for in certain sub-populations (**Supplementary Table 3**). Future work may include estimating allele frequencies for the absent pathogenic variants by incorporating the UnseenEst method that was successfully applied to estimate unseen variants in ExAC^28^.

Overall, our method provides a generalizable and robust framework to estimate disease prevalence for recessive forms of LGMD and can be adapted to estimate prevalence for other recessive diseases. By utilizing a Bayesian framework on data from the largest population reference panels (gnomAD and ExAC), this method can obtain more refined allele frequencies for rare pathogenic variants and include additional pathogenic variants from other disease databases to achieve improved disease prevalence estimates. This includes a framework for estimating the allele frequency priors, where functional annotation and CADD score groupings were used as an example. Future work will involve exploring more informative priors to improve estimates. Lastly, we have made our scripts and data available (see **Methods**), which can be easily adapted to other recessive disease genes of interest to calculate reproducible and robust estimates.

Published prevalence estimates for recessive LGMD are generally from epidemiological research studies, which are vulnerable to inaccuracies associated with delays in diagnosis or misdiagnosis, variation in diagnostic criteria used, and biases introduced by the specific population sampled^29^. By applying a Bayesian method to a genetic database, our method provides reliable disease prevalence estimates for recessive LGMD from the genetics perspective.

## Methods

### Identification of pathogenic variants

For each disease gene, variants were downloaded from the gnomAD database^30, 31^, which includes both exonic and flanking intronic variants. The Emory Genetics Laboratory (EGL) and ClinVar databases^32^ were used to annotate known pathogenic variants. Retrieved variants were first filtered based on their allele frequencies. Only variants whose minor allele frequencies are less than 0.05% in the gnomAD database were kept, unless they have been annotated as “pathogenic” or “pathogenic/likely pathogenic” in either of these two databases (EGL and ClinVar). Using the ACMG guidelines for defining pathogenic variants^11, 32^, we classified loss of function type variants as pathogenic (e.g. frameshift, stop gain, splicing donor, splicing acceptor) whether or not they were listed as pathogenic in the EGL or Clinvar databases. For the other types of variants, as long as they were annotated as pathogenic in either the EGL or ClinVar database, they were classified as pathogenic.

The above analysis is limited to known pathogenic variants and loss of function variants. We used the Combined Annotation Dependent Depletion (CADD) score^33^ cut-offs to include more variants as potentially pathogenic variants. We applied two CADD Phred-scaled score cut-offs at 20 and 30 to include variants with predicted top 1% and 0.1% deleteriousness, respectively. For further comparison, we also included all rare (AF<0.05%) missense variants to get the upper bound of estimated disease prevalence.

### Bayesian Estimation of Allele Frequencies and Disease Prevalence

The development of the disease prevalence estimator builds upon a previous published method and is detailed below.

#### Problem setting and prior assumptions

For a single variant, we would assume the observed allele count of the variant follows a binomial distribution *Binomial*(*q*_*i*_, 2*n*_*i*_), where *n*_*i*_ is the number of individuals having genotypes genotyped at this position in the database and *q*_*i*_ is the true allele frequency for this variant.

Since the conditional distribution of the observed allele count for a variant conditioned on the allele frequency *q*_*i*_ is a binomial distribution, we introduced a conjugate prior of *q*_*i*_,*q*_*i*_ ~ *Beta*(*ν*_*c:i*∈*c*_, *w*_*c:i*∈*c*_), where *ν*_*c:i*∈*c*_ and *w*_*c:i*∈*c*_ denote the prior parameters for variants belonging to the category c, which are estimated using method of moments based on all variants data provided in the ExAC database^13^. We grouped all variants into eight categories: frameshift, splice acceptor, splice donor, stop gained, missense, UTR (including 3’ and 5’ UTR), other exonic and other variants. Variants of different functional annotation are known to have different allele frequency spectrum^34^. By categorizing variants based on their functional consequence, we can get a better estimate of their allele frequencies which would be otherwise averaged among all variants, leading to an inflated allele frequency estimate for variants with more deleterious functional consequences. The allele frequencies for variants of a functional annotation are assumed to follow the same prior distribution across all genes.

In an additional analysis exploring possible more informative priors, the CADD score was incorporated in the prior as score ranges in four groups: <5, 5-30, >30 and those without a score. In combination with the 8 functional categories (mentioned above), created a total of 32 categories with allele frequency priors then calculated similarly across all genes.

#### Posterior distribution of allele frequencies

The posterior distribution of the allele frequency *q*_*i*_ given the observed allele counts *x*_*i*_ and prior assumption on the allele frequency would be

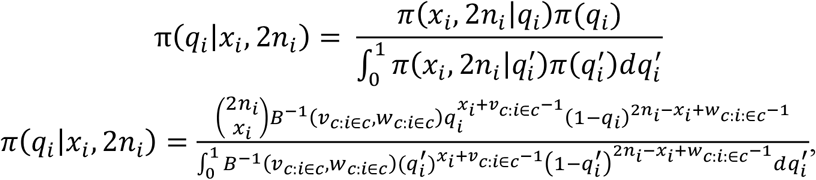

Where *B*^−1^(*ν*_*c:i*∈*c*_, *w*_*c:i*∈*c*_) is the inverse of the beta function *B*(*ν*_*c:i*∈*c*_, *w*_*c:i*∈*c*_) which makes the total probability of beta distribution *Beta*(*ν*_*c:i*∈*c*_, *w*_*c:i*∈*c*_) be 1. Based on the equation above, we can infer that the posterior distribution of *q*_*i*_ is a beta distribution: *Beta*(*x*_*i*_ + *ν*_*c:i*∈*c*_, 2*n*_*i*_ − *x*_*i*_ + *w*_*c:i*∈*c*_). For pathogenic variants (from EGL and ClinVar) unseen in the population reference panel, we would take *x*_*i*_ being 0 and *n*_*i*_ being the corresponding sample size in the mixed population or the specific sub population correspondingly.

#### Posterior estimation of disease prevalence

For monogenic rare diseases the disease prevalence would be *D* = [1 − Π_*i*_(1 − *q*_*i*_)]^2^. It is the probability of both two copies of the disease gene having at least one pathogenic variant. Here, we assume that effect for each pathogenic variant in the disease gene is the same and independent assortment of pathogenic variants. In practice, *q*_*i*_’s that we are using here are quite small, therefore, we can use *D* ≈ (Σ_*i*_*q*_*i*_)^2^ to approximate the disease prevalence.

From the equation listed above, the approximation of prevalence involves the sum of multiple variables from different beta distributions, since *q*_*i*_ follows a beta distribution. Empirically, we use a normal distribution to approximate the distribution of the sum of beta variables^35^.

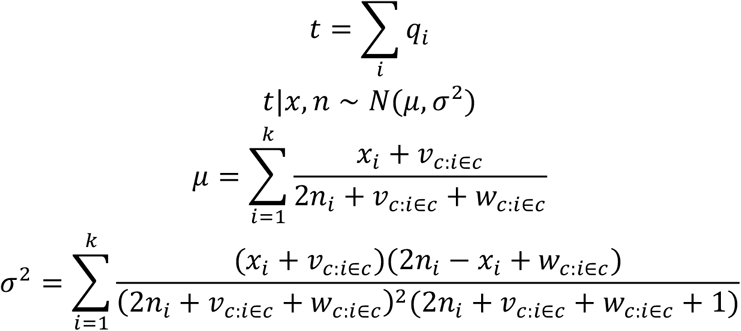

Therefore, the posterior of the prevalence would follows a chi-square distribution with the degree of freedom being 1 and a non-central parameter. Using 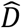 to denote the approximation term for disease prevalence (Σ*q*_*i*_)^2^, we can get 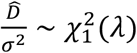 and 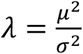. With the posterior distribution of the approximated disease prevalence in this form, it is easy for us to get the prevalence estimator and its confidence intervals. We are using the expectation (*λ* + 1)*σ*^2^ of the distribution as the prevalence estimator here. The lower bound of the estimator with the confidence 1 − *α* would be 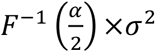, where *F*(.) is the cumulative distribution function for the chi-square distribution. Similarly, the upper bound for the estimator would be 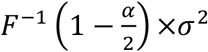. We are using *α* = 0.05 here to get the 95% confidence interval for our prevalence estimators.

#### Direct estimation of disease prevalence in genetic databases

For comparison, we also estimated disease prevalence by using the observed allele frequency of a pathogenic variant in genetic databases as the direct estimator for *q*_*i*_ (without beta prior). More specifically, the disease prevalence can be estimated by:

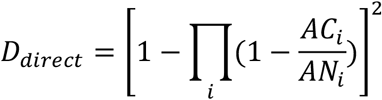

where *AC*_*i*_ is the allele count for the variant *i* and *AN*_*i*_ is the corresponding allele number in the position. As above, for a given disease or a subtype, the product is taken over all identified pathogenic variants in the disease gene, where *i* is the index of those identified pathogenic variants.

The scripts for estimating recessive disease prevalence based on our Bayesian framework and also direct calculation are available at https://github.com/leklab/prevalence_estimation.

### Used URLs

gnomAD: http://gnomad.broadinstitute.org/downloads;

EGL genetics database: http://www.egl-eurofins.com/emvclass/emvclass.php;

ClinVar database (the version used here is 20180429, may be updated now): ftp://ftp.ncbi.nlm.nih.gov/pub/clinvar/vcf_GRCh37/

ExAC database: ftp://ftp.broadinstitute.org/pub/ExAC_release/release1/manuscript_data/;

BRAVO database: https://bravo.sph.umich.edu/freeze5/hg38/

## Acknowledgements

N.J.L. is the recipient of a NHMRC CJ Martin Early Career Fellowship and an American Australian Association scholarship. S.P. was supported by the Estonian Research Council grant (PUTJD827).

## Author contributions

W.L. and M.L. conceived the study and developed the statistical framework

W.L. implemented the method

W.L., S.P., N.J.L. and G.Z. conducted prevalence estimation

N.I., P.M., N.E.J., C.C.W., B.A.W., D.E.A., L.E.R. analyzed genetic analysis and compared epidemiology results

W.L., S.P., N.J.L. and M.L. wrote the manuscript

All authors read and approved the manuscript.

## Supplementary Tables

**Supplementary Table 1:** Gene lengths and numbers of pathogenic variants used for prevalence estimation

| Subtype/Gene | Gene length (Kb) | Coding sequence length (bp) Transcript ID | # pathogenic variants used in our method | # published causative variants<sup>44</sup> | # variants in ClinVar or EGL but not in gnomAD |
| --- | --- | --- | --- | --- | --- |
| 2A/<br>CAPN3 | 29.2 | 2,466<br>NM_000070 | 149(59 annotated in ClinVar/EGL) | 289 | 40 |
| 2B/<br>DYSF | 80.4 | 6,243<br>NM_003494 | 225<br>(83 annotated in ClinVar/EGL) | 213 | 93 |
| 2C/<br>SGCG | 1.6 | 876<br>NM_000231 | 34<br>(8 annotated in ClinVar/EGL) | 17 | 6 |
| 2D/<br>SGCA | 9.0 | 1,164<br>NM_000023 | 51<br>(11 annotated in ClinVar/EGL) | 63 | 14 |
| 2E/<br>SGCB | 5.7 | 957<br>NM_000232 | 35<br>(7 annotated in ClinVar/EGL) | 31 | 8 |
| 2F/<br>SGCD | 13.5 | 873<br>NM_000337 | 26<br>(1 annotated in ClinVar/EGL) | 8 | 6 |
| 2G/<br>TCAP | 2.7 | 504<br>NM_003673 | 20<br>(1 annotated in ClinVar/EGL) | 3 | 4 |
| 2I/<br>FKRP | 19.7 | 1,488<br>NM_024301 | 49<br>(16 annotated in ClinVar/EGL) | 95 | 20 |
| 2L/<br>ANO5 | 14.0 | 2,742<br>NM_213599 | 109<br>(32 annotated in ClinVar/EGL) | 29 | 15 |

The table indicates numbers of pathogenic variants used for estimating prevalence in our method, compared with numbers of pathogenic variants obtained from a recent publication, which gathered variants from Leiden Open Variation Database and the Human Gene Mutation Database for each LGMD2 subtype.

**Supplementary Table 2:**
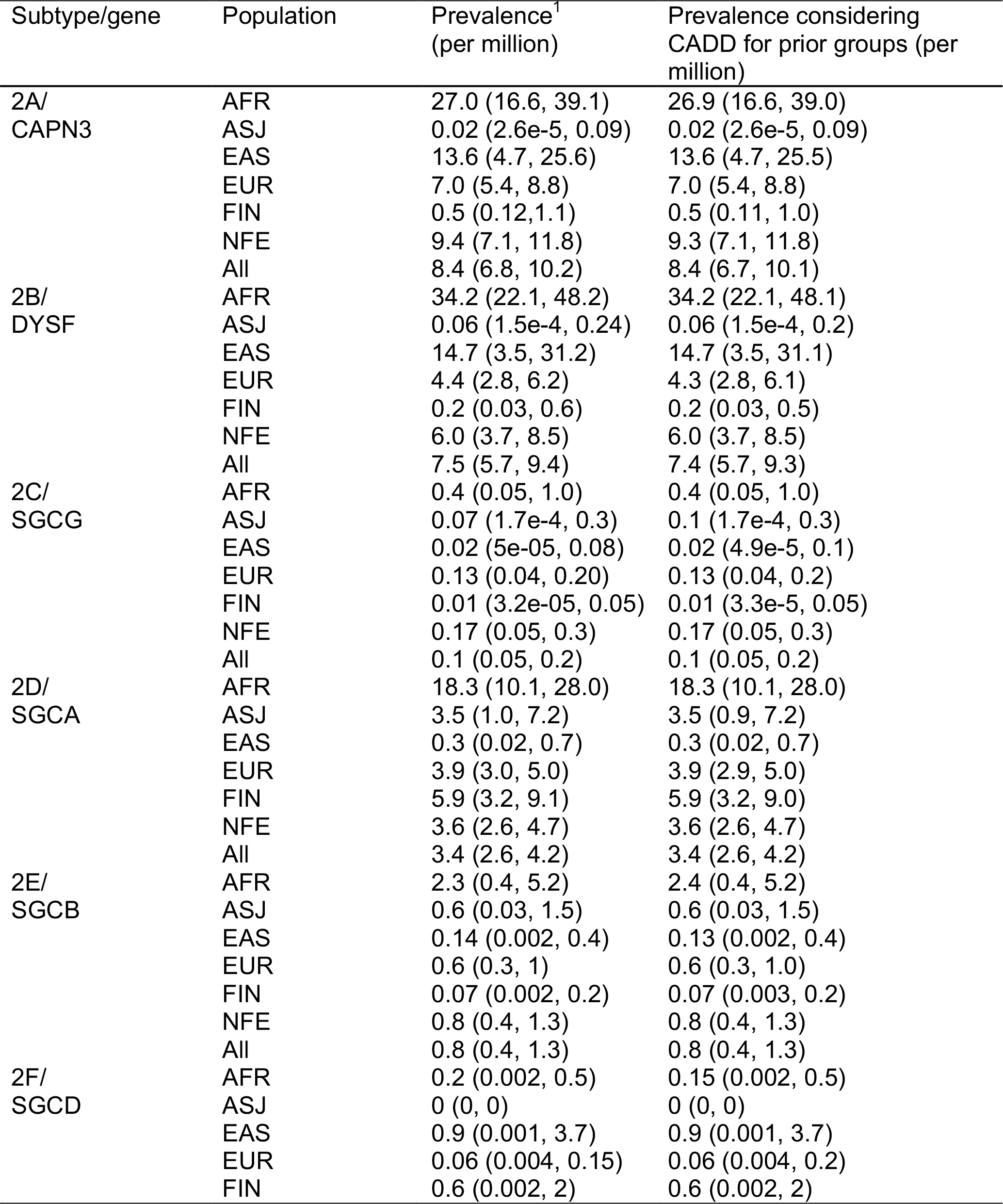
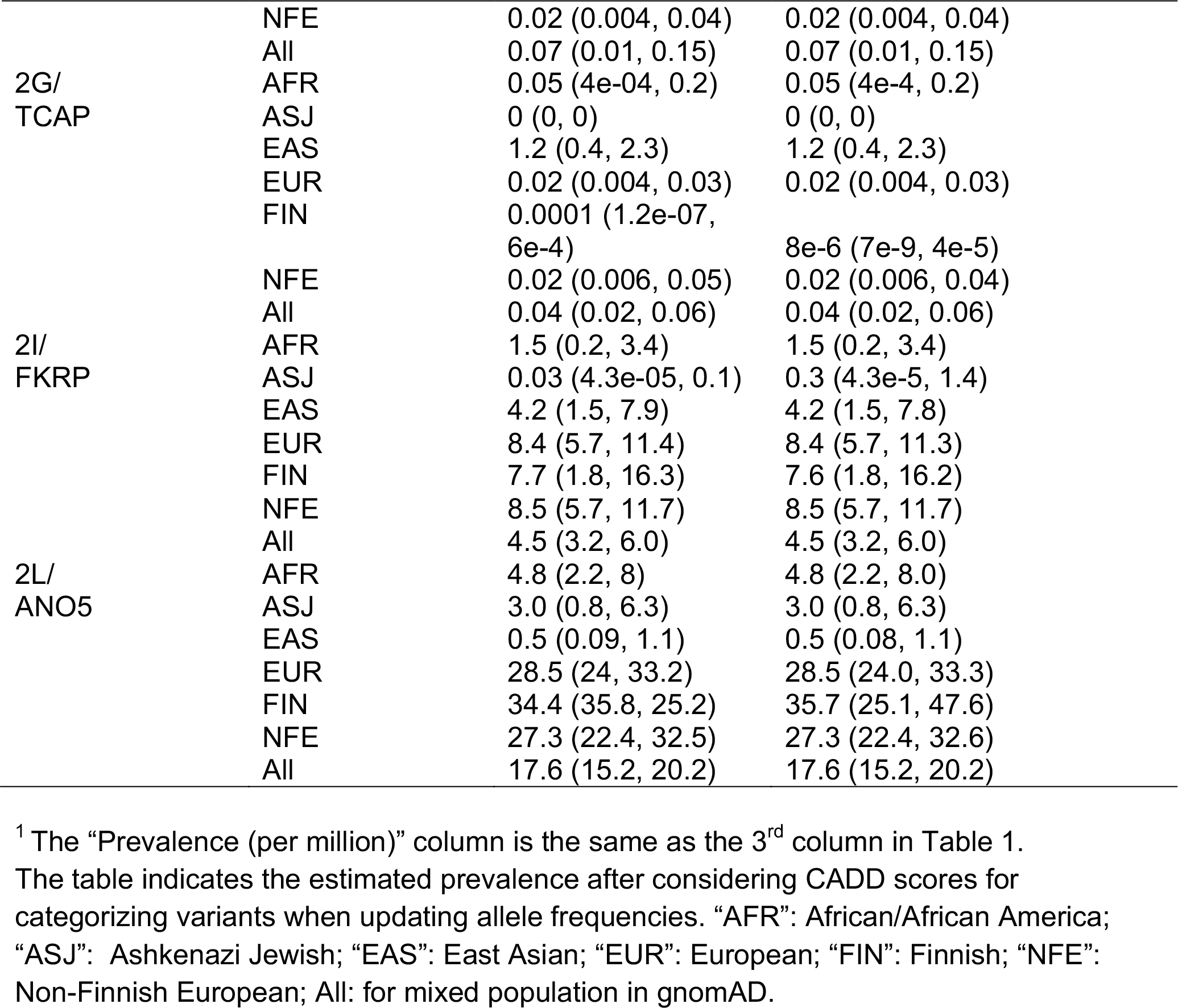
Prevalence results considering CADD scores when estimating categorized priors for allele frequencies

**Supplementary Table 3:**
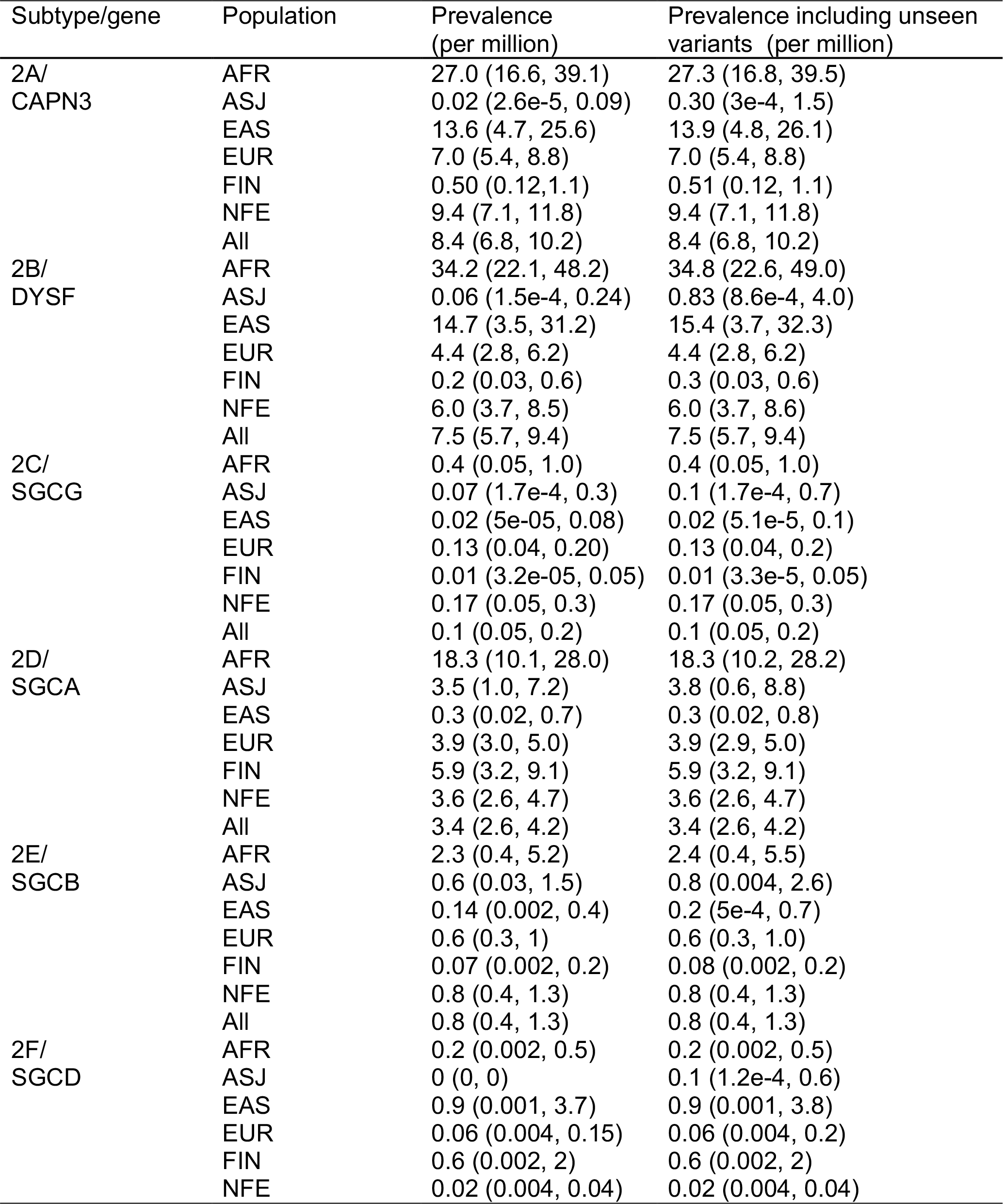
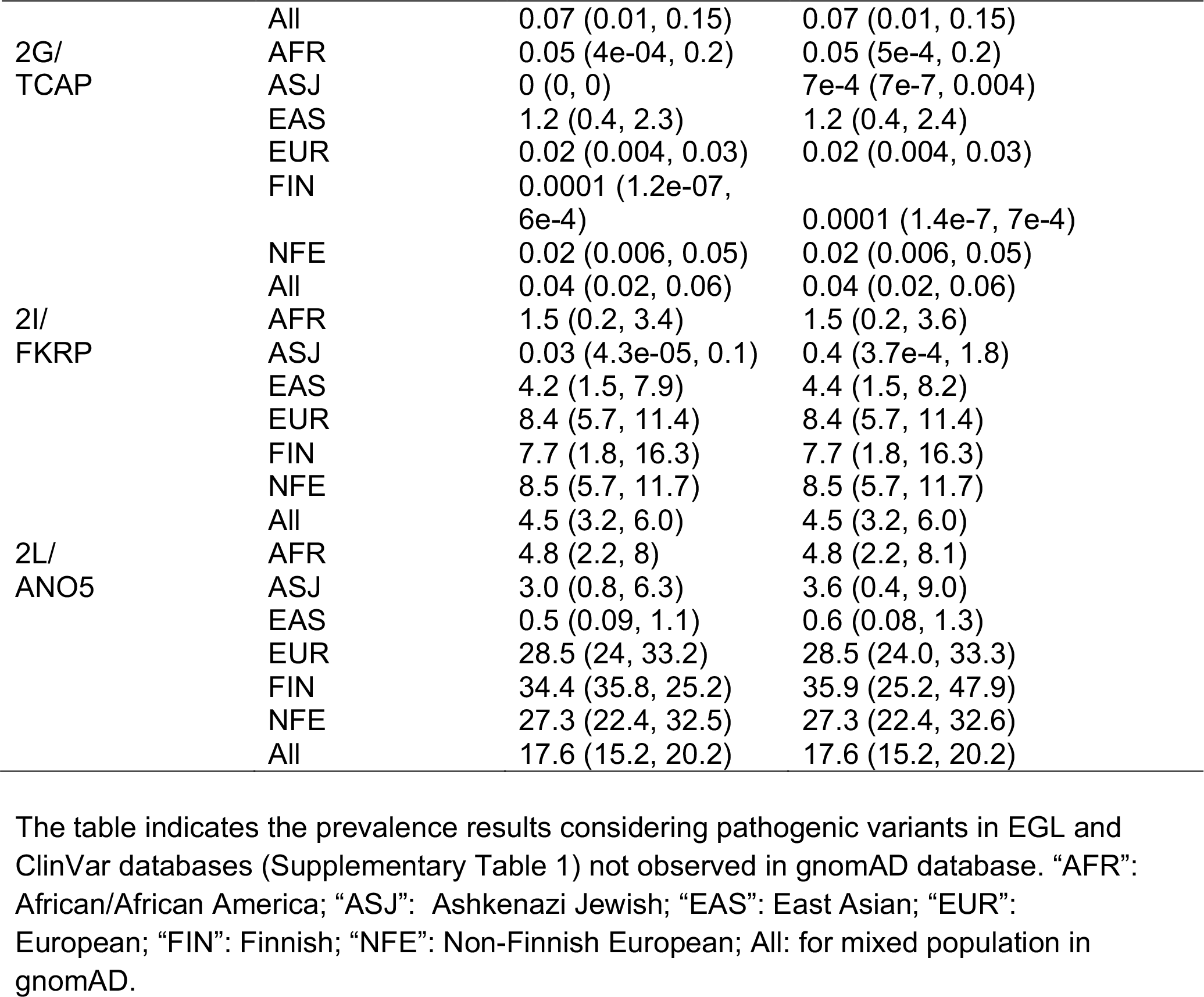
Prevalence estimates (per million) including reported pathogenic variants unseen in gnomAD

